# The Role of Glucocorticoid and Nicotinic Acetylcholine Receptors in the Reward-Enhancing Effects of Nicotine in the ICSS Procedure in Male and Female Rats

**DOI:** 10.1101/2024.07.03.601962

**Authors:** Ranjithkumar Chellian, Azin Behnood-Rod, Adriaan W. Bruijnzeel

## Abstract

Tobacco use disorder is a chronic disorder that affects more than one billion people worldwide and causes the death of millions each year. The rewarding properties of nicotine are critical for the initiation of smoking. Previous research has shown that the activation of glucocorticoid receptors (GRs) plays a role in nicotine self-administration in rats. However, the role of GRs in the acute rewarding effects of nicotine are unknown. In this study, we investigated the effects of the GR antagonist mifepristone and the nicotinic acetylcholine receptor (nAChR) antagonist mecamylamine on the reward-enhancing effects of nicotine using the intracranial self-stimulation (ICSS) procedure in adult male and female rats. The rats were prepared with ICSS electrodes in the medial forebrain bundle and then trained on the ICSS procedure. Nicotine lowered the brain reward thresholds and decreased response latencies similarly in male and female rats. These findings suggest that nicotine enhances the rewarding effects of ICSS and has stimulant properties. Treatment with the GR antagonist mifepristone did not affect the reward-enhancing effects of nicotine but increased response latencies, suggesting a sedative effect. Mecamylamine did not affect the brain reward thresholds or response latencies of the control rats, but prevented the nicotine-induced decrease in brain reward thresholds and reward latencies. These findings indicate that the rewarding effects of nicotine are mediated via the activation of nAChRs, and that the activation of GRs does not contribute to the acute rewarding effects of nicotine. These studies enhance our understanding of the neurobiological mechanisms underlying tobacco use disorder.

## Introduction

Tobacco use disorder is the leading preventable cause of death worldwide. Worldwide, there are about 1.3 billion people who smoke and this leads to the death of over 8 million people annually ^1^. Smoking increases the risk for a wide range of diseases, including chronic obstructive pulmonary disease, cardiovascular disease, and cancer ^2-5^. Despite extensive public health campaigns and the availability of a wide range of smoking cessation aids, the smoking rate remains high ^6^. Furthermore, relapse is common among people who try to quit smoking ^7^. A better understanding of the neurobiological mechanisms underlying nicotine addiction is critical for developing more effective treatments for smoking cessation.

Nicotine induces feelings of mild euphoria (e.g., a nicotine buzz) in smokers and vapers ^8,9^. The rewarding effects of nicotine play a role in the initiation and maintenance of smoking and vaping ^10^. Nicotine mediates its rewarding effects at least partly by activating α4β2* and α6β2* containing nicotinic acetylcholine receptors (nAChRs) on dopamine neurons in the VTA ^11^. This subsequently leads to the release of dopamine in the nucleus accumbens, and activation of dopamine D1 receptors. The activation of dopamine D1 receptors contributes to the rewarding effects of nicotine and thereby promoting the continuation of nicotine use ^12,13^.

Exposure to stressors increases smoking and impairs the ability to resist smoking ^14,15^. People who are stressed may smoke more to cope with stress or to alleviate their feelings of tension and anxiety ^16^. Glucocorticoids play a critical role in the reinforcing effects of drugs, as indicated by the fact that adrenalectomy decreases the acquisition of drug intake and reduces drug self-administration ^17,18^. Additionally, both adrenalectomy and corticosterone synthesis inhibition have been found to prevent stress-induced sensitization of the motor effects of psychostimulants ^19,20^. The glucocorticoid receptor (GR, Ki 0.09 nM) and progesterone receptor (Ki 1 nM) antagonist mifepristone (also referred to as RU 486 or RU 38486) has been widely used to investigate the role of glucocorticoids in drug intake ^21-25^. Mifepristone prevents drug-induced sensitization of locomotor responses, indicating that GRs play a critical role in drug-induced neuronal adaptations ^26^. Furthermore, mifepristone decreases the self-administration of nicotine and other stimulants such as cocaine and amphetamine in rats ^27-29^.

It is currently not known if GR blockade diminishes nicotine intake by decreasing the acute rewarding properties of nicotine. Therefore, in this study, we aim to elucidate the effects of GR blockade on the reinforcing properties of nicotine using the intracranial self-stimulation (ICSS) procedure in male and female Wistar rats. The ICSS procedure has been widely used to investigate the effects of drugs of abuse on brain reward function ^30,31^. A decrease in brain reward thresholds indicates that rats are more sensitive to rewarding electrical stimuli and reflects a potentiation of brain reward function. Furthermore, at the end of the study, we investigated whether mecamylamine affects the reward-enhancing effects of nicotine in the ICSS procedure. Mecamylamine is a non-selective and non-competitive antagonist of nAChRs and is most potent at inhibiting α3β4 and less potent at inhibiting α4β2, α3β2, and α7 nAChRs ^32^. Although mecamylamine has been shown to decrease nicotine self-administration ^33^, its effect on the reward-enhancing effects of noncontingently administered nicotine in males and females has not been investigated. Therefore, we also investigated whether mecamylamine blocks the acute reward-enhancing and stimulatory effects of nicotine in male and female rats.

## 2. Materials and Methods

### 2.1. Animals

Adult male (200–250 g, 8-9 weeks of age; N=11) and female (175–225 g, 8-9 weeks of age; N=10) Wistar rats were purchased from Charles River (Raleigh, NC). The rats were housed with a rat of the same sex in a climate-controlled vivarium on a reversed 12 h light-dark cycle (light off at 7 AM). The rats were singly housed after the implantation of the ICSS electrodes. Food and water were available ad libitum in the home cage. The experimental protocols were approved by the University of Florida Institutional Animal Care and Use Committee (IACUC). All experiments were performed in accordance with relevant IACUC guidelines and regulations and in compliance with ARRIVE guidelines 2.0 (Animal Research: Reporting of *In Vivo* Experiments).

### 2.2. Drugs

Mifepristone (Cayman chemical company, Ann Arbor, MI, USA) was dissolved in 10% dimethylformamide (Sigma-Aldrich, St. Louis, MO, USA), 10% kolliphor (Sigma-Aldrich, St. Louis, MO, USA) and mixed in sterile saline. (-)-Nicotine hydrogen tartrate (NIDA Drug Supply Program) and mecamylamine hydrochloride (NIDA Drug Supply Program) were dissolved in sterile saline. Mifepristone was administered intraperitoneally (IP), while mecamylamine and nicotine were administered subcutaneously (SC) in a volume of 1 ml/kg body weight. The nicotine dose is expressed as base and the mecamylamine dose is expressed as salt.

### 2.3. Mifepristone and nicotine treatment

Mifepristone (0, 3, 10, and 30 mg/kg, IP) and nicotine (0 and 0.3 mg/kg, SC) were administered according to a Latin square design. Mifepristone was administered 90 min before the ICSS sessions and nicotine was administered 15 min before the ICSS sessions. There were at least 72 h between treatment days, at least one ICSS sessions was conducted without any drug treatment before treatment days. The doses of mifepristone were based on a previous study that showed that these doses decrease nicotine intake in rats ^34^.

### 2.4. Mecamylamine and nicotine treatment

Mecamylamine (0 and 3 mg/kg, SC) and nicotine (0 and 0.3 mg/kg, SC) were administered according to a Latin square design. Mecamylamine was administered 30 min before the ICSS sessions and nicotine was administered 15 min before the ICSS sessions. Twenty-four hours after treatment, ICSS sessions were conducted without any drug treatment. There were at least 48 h between treatment days. The mecamylamine dose was based on our previous study that showed that 3 mg/kg of mecamylamine diminishes the nicotine self-administration induced decrease in brain reward thresholds and latencies in rats ^35^.

### 2.5. Electrode implantation and ICSS procedure

Adult male (n=11) and female (n=10) rats were prepared with ICSS electrodes in the medial forebrain bundle and trained on the ICSS procedure ^35-37^. Briefly, the rats were anesthetized with an isoflurane and oxygen vapor mixture and placed in a stereotaxic frame (David Kopf Instruments, Tujunga, CA, USA). Electrodes were implanted in the medial forebrain bundle with the incisor bar set 5 mm above the interaural line (anterior-posterior -0.5 mm, medial-lateral ±1.7 mm, dorsal-ventral -8.3 mm from dura). The rats were trained on a modified discrete-trial ICSS procedure in operant conditioning chambers (Med Associates, Georgia, VT, USA), housed within sound-attenuating enclosures. A five cm wide metal response wheel was centered on a sidewall, with a photobeam detector recording every 90 degrees of rotation. Brain stimulation was delivered by constant current stimulators (Model 1200C, Stimtek, Acton, MA, USA). The rats were trained to turn the wheel on a fixed ratio 1 (FR1) schedule of reinforcement. Each quarter-turn of the wheel resulted in the delivery of a 0.5-second train of 0.1 ms cathodal square-wave pulses at a frequency of 100 Hz. After acquiring responding for stimulation on the FR1 schedule (100 reinforcements within 10 minutes), the rats were trained on a discrete-trial current-threshold procedure. The discrete-trial current-threshold procedure is a modification of a task developed by Kornetsky and Esposito ^38^, and previously described in detail ^35-37^. Each trial began with the delivery of a noncontingent electrical stimulus, followed by a 7.5-s response window during which the animals could respond for a second identical stimulus. A response during this 7.5-s window was labeled a positive response, and the lack of a response was labeled a negative response. During the 2-s period immediately after a positive response, additional responses had no consequences. The inter-trial interval (ITI), which followed a positive response or the end of the response window, had an average duration of 10 s (ranging from 7.5 to 12.5 s). Responses during the ITI resulted in a 12.5-s delay of the onset of the next trial. During the training sessions, the duration of the ITI and delay periods induced by time-out responses were increased until the animals performed consistently at standard test parameters. The training was completed when the animals responded correctly to more than 90% of the noncontingent electrical stimuli. It took 2-3 weeks of training for most rats to meet this response criterion. The rats were then tested on the current-threshold procedure in which stimulation intensities varied according to the classical psychophysical method of limits. Each test session consisted of four alternating series of descending and ascending current intensities starting with a descending sequence. Blocks of three trials were presented to the rats at a given stimulation intensity, and the intensity was altered between blocks of trials by 5 µA steps. The initial stimulus intensity was set 30 µA above the baseline current threshold for each animal. Each test session typically lasted 30-40 min and provided two dependent variables for behavioral assessment (brain reward thresholds and response latencies). The brain reward threshold (µA) was defined as the midpoint between stimulation intensities that supported responding and stimulation intensities that failed to support responding. The response latency (s) was defined as the time interval between the beginning of the noncontingent stimulus and a response. When the brain reward thresholds were stable (less than 10% variation within a 5-day period), the rats were treated with drugs before the ICSS sessions. A decrease in reward thresholds is indicative of an increase in reward function ^38^. Drugs with sedative effects increase the response latencies, and stimulants decrease the response latency ^31,39^.

### 2.6. Statistics

The brain reward thresholds and the response latencies were expressed as a percentage of the 3-day baseline that was obtained prior to the drug treatments. The percentage changes in thresholds and latencies were analyzed with three-way ANOVAs, with nicotine and mifepristone or mecamylamine treatment as within-subject factors, and sex as a between-subjects factor. For all statistical analyses, significant interaction effects found in the ANOVAs were followed by Bonferroni’s post hoc tests to determine which groups differed. P-values less than or equal to 0.05 were considered significant. Data were analyzed with IBM SPSS Statistics version 29 and GraphPad Prism version 10.1.2. The figures were generated using GraphPad Prism version 10.1.2.

## 3. Results

### Effects of mifepristone on the rewarding effect of nicotine in the ICSS procedure

#### Reward thresholds

Treatment with nicotine lowered the brain reward thresholds and this effect was not affected by the sex of the rats (Fig. 1A, Nicotine treatment: F1,19=27.256, P < 0.001; Sex: F1,19=0.212, NS; Nicotine treatment × Sex: F3,57=0, NS). Treatment with mifepristone did not affect the reward thresholds (Fig. 1A, Mifepristone treatment: F3,57=1.57, NS; Mifepristone treatment × Nicotine treatment: F3,57=0.243, NS; Mifepristone treatment × Sex: F3,57=0.663, NS; Mifepristone treatment × Nicotine treatment × Sex: F3,57=0.365, NS).

**Figure 1.**
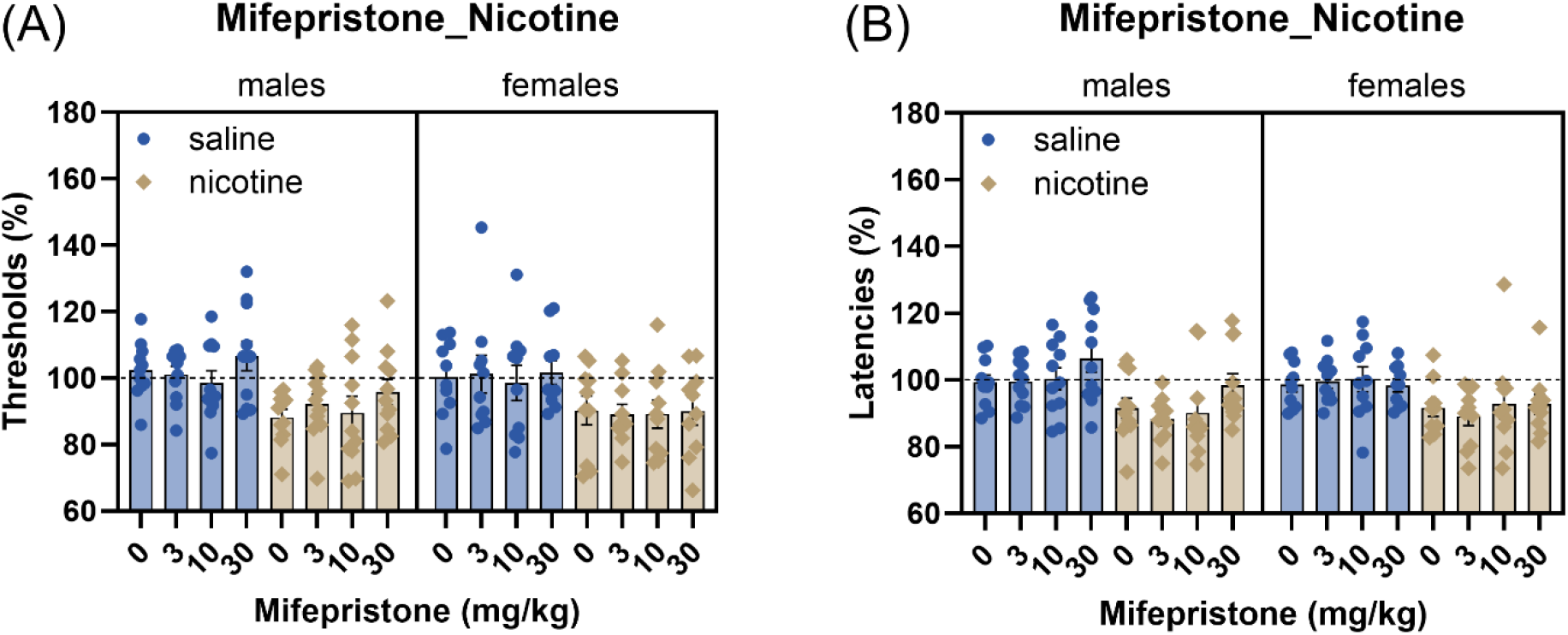
Mifepristone did not affect the reward-enhancing and stimulant effects of nicotine. Nicotine (0.3 mg/kg, sc) decreased the brain reward thresholds (A) and the response latencies (B) in the males and the females. Male (N=11); female (N=10). Data are expressed as means ± SEM.

#### Response latencies

Treatment with nicotine also decreased the response latencies and this was not affected by the sex of the rats (Fig. 1B, Nicotine treatment: F1,19=29.097, P < 0.001; Sex: F1,19=0.228, NS; Nicotine treatment × Sex: F3,57=0.305, NS). Treatment with mifepristone increased the latencies and this was not affected by treatment with nicotine or the sex of the rats (Fig. 1B, Mifepristone treatment: F3,57=2.919, P < 0.05; Mifepristone treatment × Nicotine treatment: F3,57=0.621, NS; Mifepristone treatment × Sex: F3,57=2.452, NS; Mifepristone treatment × Nicotine treatment × Sex: F3,57=0.065, NS)

### Effects of mecamylamine on the rewarding effect of nicotine in the ICSS procedure

#### Reward thresholds

Treatment with nicotine lowered the brain reward thresholds and this was not affected by the sex of the rats (Fig. 2A, Nicotine treatment: F1,19=17.494, P < 0.001; Sex: F1,19=1.147, NS; Nicotine treatment × Sex: F1,19=0.579, NS). Treatment with mecamylamine prevented the nicotine-induced decrease in brain reward thresholds (Fig. 2A, Mecamylamine treatment: F1,19=20.246, P < 0.001; Mecamylamine treatment × Nicotine treatment: F1,19=6.399, P < 0.05). The effect of mecamylamine on the brain reward thresholds was not affected by the sex of the rats (Mecamylamine treatment × Sex: F1,19=1.877, NS; Mecamylamine treatment × Nicotine treatment × Sex: F1,19=0.858, NS). Since there was no significant Mecamylamine treatment × Nicotine treatment × Sex interaction, the post hoc test was conducted on a group that included both males and females. The post hoc analysis showed that mecamylamine prevented the nicotine-induced decrease in brain reward thresholds in the combined group of males and females (Fig. 2B).

**Figure 2.**
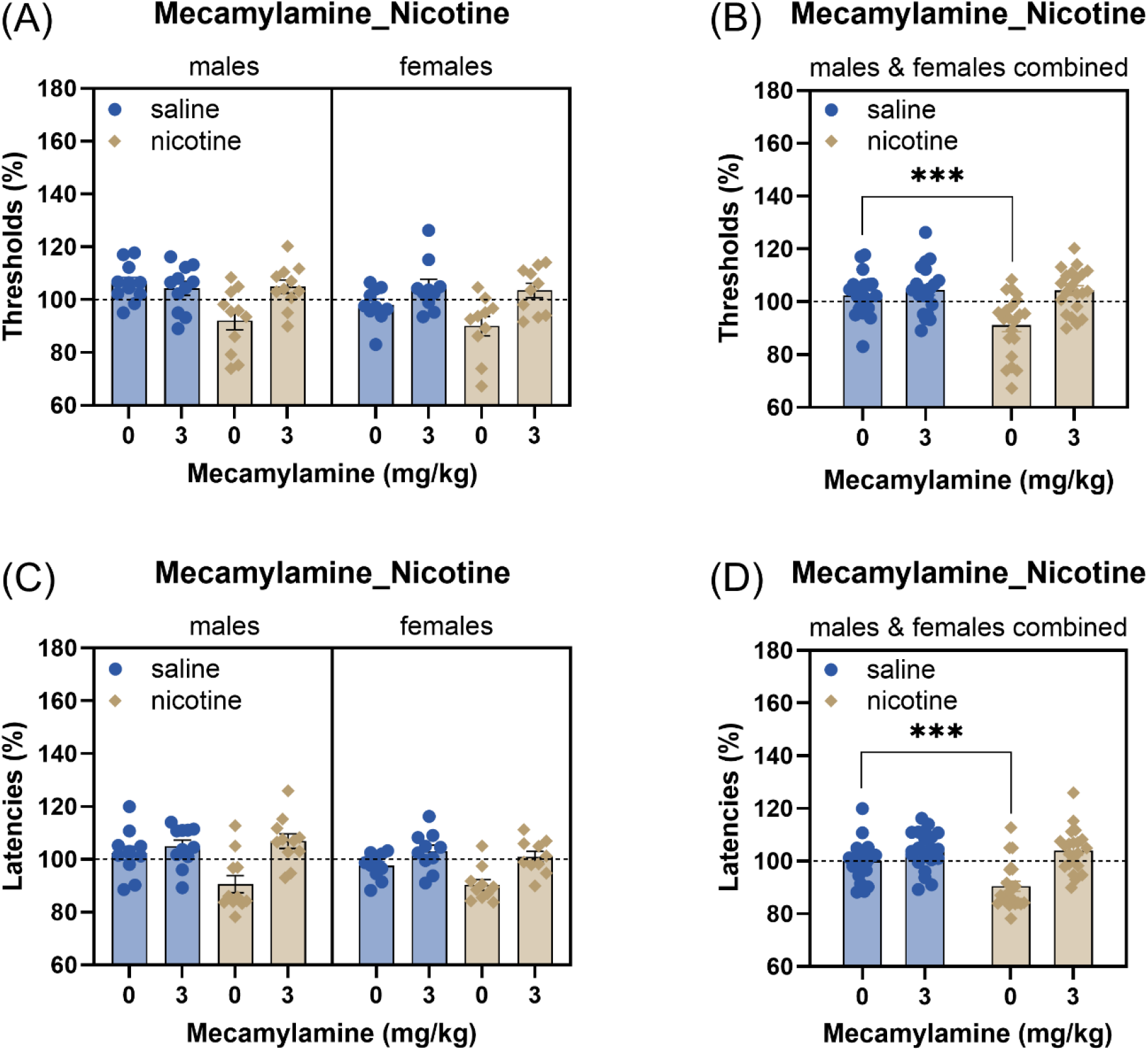
Mecamylamine prevented the reward-enhancing and stimulant effects of nicotine. Nicotine (0.3 mg/kg, sc) decreased the brain reward thresholds (A, B) and the response latencies (C, D) in both males and females, with data shown separately and combined for both sexes. Mecamylamine prevented the decrease in reward thresholds and latencies (B, D). Male (N=11); female (N=10). Asterisks indicate lower brain reward thresholds or shorter latencies compared to rats treated with saline and vehicle. *** P<0.001. Data are expressed as means ± SEM.

#### Response latencies

Treatment with nicotine decreased the response latencies and this was not affected by the sex of the rats (Fig. 2C, Nicotine treatment: F1,19=12.184 P < 0.01; Sex: F1,19=2.322, NS; Nicotine treatment × Sex: F1,19=0.01, NS.). Mecamylamine prevented the nicotine induced decrease in the response latencies (Fig. 2C, Mecamylamine treatment: F1,19=32.175, P < 0.001; Mecamylamine treatment × Nicotine treatment: F1,19=6.878, P < 0.05). The effect of mecamylamine on the response latencies was not affected by the sex of the rats (Mecamylamine treatment × Sex: F1,19=0.2, NS; Mifepristone treatment × Nicotine treatment × Sex: F1,19=1.391, NS). The posthoc showed that mecamylamine prevented the nicotine-induced decrease in response latencies in the combined group of males and females (Fig. 2D).

## Discussion

In this study, we investigated the effects of GR and nAChR blockade on the reward-enhancing effects of nicotine in the ICSS procedure in male and female Wistar rats. Our studies showed that nicotine lowered the brain reward thresholds and decreased the response latencies. The effects of nicotine on the reward thresholds and response latencies was not affected by the sex of the rats. Treatment with mifepristone did not affect the reward-enhancing effects of nicotine. However, mifepristone increased the response latencies, suggesting a sedative effect. Treatment with mecamylamine prevented the reward-enhancing and stimulant effects of nicotine. These findings indicate that the effects of nicotine, mifepristone, and mecamylamine are not affected by the sex of the rats. Additionally, our results indicate that blockade of nAChRs, but not GRs, diminishes the acute rewarding effects of nicotine.

The administration of nicotine lowered the brain reward thresholds, indicating that nicotine potentiates brain reward function. This finding is consistent with other studies showing that nicotine enhances the rewarding effects of ICSS and induces conditioned place preference ^31,40,41^. In our study, there was no sex difference in the effects of nicotine on brain reward function. Similarly, a prior study also found no sex difference in the reward-enhancing effects of nicotine in the ICSS procedure ^36^. In the current study, we used only one dose of nicotine, but prior research showed that there is also no sex difference in the rewarding enhancing effects of lower doses of nicotine in the ICSS procedure ^36^. However, we are not aware of any studies that have investigated sex differences in the effects of higher doses of nicotine in the ICSS procedure. Nicotine also induces conditioned place preference in both male and female rodents, but higher doses of nicotine are required to induce conditioned place preference in females 42,43.

One of the goals of the present study was to investigate the role of GRs in the acute rewarding effects of nicotine. Treatment with the GR antagonist mifepristone did not affect the brain reward thresholds in the saline controls and did not prevent the nicotine-induced decrease in brain reward thresholds. These findings suggest that GR blockade with mifepristone does not affect the acute reward-enhancing effects of nicotine as measured by ICSS. In one of our previous studies, we found that mifepristone decreases the self-administration of nicotine in rats ^29^. The current findings do not support a role for GRs in the acute rewarding properties of nicotine. Therefore, GRs may affect nicotine intake via mechanisms independent of inhibiting the acute rewarding effects of nicotine. Although the present study showed that GRs do not play a role in the acute rewarding effects of nicotine, previous research has shown that GR activation is essential for the development of behavioral sensitization to nicotine ^44^.

Interestingly, while mifepristone did not affect the reward thresholds, it did increase response latencies. This increase in response latencies was not affected by nicotine treatment or the sex of the rats. The differential effects of mifepristone on reward thresholds and response latencies suggest that GR blockade may induce some sedation without affecting brain reward function. In a prior study, we also found that treatment with mifepristone (30 mg/kg) decreases locomotor activity and rearing in the open field test, further suggesting that mifepristone has some sedative effects ^29^.

In contrast to mifepristone, the nAChR antagonist mecamylamine significantly affected nicotine-induced changes in both reward thresholds and response latencies. Mecamylamine prevented the nicotine-induced decrease in brain reward thresholds and response latencies in a group of males and females. These results are consistent with a previous study that showed that that nicotinic receptor antagonist dihydro-beta-erythroidine hydrobromide (DHβE) diminishes nicotine’s reward-enhancing effects in the ICSS procedure ^45^. Furthermore, in a previous study we found that mecamylamine prevented the nicotine self-administration induced decrease in brain rewards thresholds and response latencies ^35^. Taken together, these findings indicate that nAChR blockade diminishes the rewarding effects of both noncontingent nicotine administration and nicotine self-administration.

In conclusion, our study demonstrates that nAChR blockade with mecamylamine reduces the rewarding effects of nicotine in both male and female rats, without inducing sedative effect. In contrast, GR blockade with mifepristone did not diminish the rewarding effects of nicotine, but did have some sedative effects. These findings suggest that the rewarding effects of nicotine are mediated via the activation of nAChRs, and that the activation of GRs does not contribute to the acute rewarding effects of nicotine. This work contributes to our understanding of the neurobiological mechanisms underlying nicotine addiction and may aid in the development of therapies for smoking cessation.

## CRediT authorship contribution statement

**R. Chellian:** Formal analysis, Investigation, Writing - Review & Editing, Visualization. **A. Behnood-Rod:** Investigation, Project administration. **A. Bruijnzeel:** Conceptualization, Formal analysis, Writing - Original Draft, Visualization, Supervision, Project administration, Funding acquisition.

